# Duplicate transcription factors *GT1* and *VRS1* regulate branching and fertile flower number in maize and *Brachypodium distachyon*

**DOI:** 10.1101/2023.03.15.532786

**Authors:** Joseph P. Gallagher, Jarrett Man, Adriana Chiaramida, Isabella Rozza, Erin L. Patterson, Morgan Powell, Amanda Schrager-Lavelle, Dilbag S. Multani, Robert Meeley, Madelaine E. Bartlett

**Affiliations:** Biology Department, University of Massachusetts, Amherst, MA; Forage Seed and Cereal Research Unit, USDA-ARS, Corvallis, OR; Corteva Agriscience, Johnston, IA

**Keywords:** HD-ZIP, CRISPR-Cas9 gene editing, inflorescence development, flower development, *de novo* domestication

## Abstract

Crop engineering and *de novo* domestication using genome editing are new frontiers in agriculture. However, outside of well-studied crops and model systems, prioritizing engineering targets remains challenging. Evolution can serve as our guide, revealing high-priority genes with deeply conserved roles. Indeed, *GRASSY TILLERS1* (*GT1*), *SIX-ROWED SPIKE1* (*VRS1*), and their homologs have repeatedly been targets of selection in domestication and evolution. This repeated selection may be because these genes have an ancient, conserved role in regulating growth repression. To test this, we determined the roles of *GT1* and *VRS1* homologs in maize (*Zea mays*) and the distantly related grass brachypodium (*Brachypodium distachyon*) using CRISPR-Cas9 gene editing and mutant analysis. *GT1* and *VRS1* have roles in floral development in maize and barley, respectively. Grass flowers are borne in branching structures called spikelets. In maize spikelets, carpels are suppressed in half of all initiated ear flowers. These spikelets can only produce single grains. We show that *gt1; vrs1-like1* (*vrl1*) mutants have derepressed carpels in ear flowers. Importantly, these plants can produce two grains per spikelet. In brachypodium, *bdgt1; bdvrl1* mutants have more branches, spikelets, and flowers than wildtype plants, indicating conserved roles for *GT1* and *VRS1* homologs in growth suppression. Indeed, maize *GT1* can suppress growth in *Arabidopsis thaliana*, separated from the grasses by *ca*. 160 million years of evolution. Thus, *GT1* and *VRS1* maintain their potency as growth regulators across vast timescales and in distinct developmental contexts. Modulating the activity of these and other conserved genes may be critical in crop engineering.

## Introduction

Plant development directly impacts crop yield. An increase in the number of fertile flowers, for example, increases the number of fruits or grains that can be produced. Improvements in fertility have been critical in the domestication and improvement of several cereal crops (Dong et al., 2019). A key to fertility is the proper development of inflorescence and floral structures.

Yield can be improved by regulating growth of reproductive structures. In the grasses, a key reproductive structure influencing yield is the spikelet. The spikelet houses the flowers within leaf-like bracts called glumes. In barley (*Hordeum vulgare*), modifying spikelet development was key in its domestication (Komatsuda et al., 2007). A triple spikelet, containing a central spikelet and two lateral spikelets, forms on each node of the inflorescence main axis, the rachis. Within each spikelet, the flower develops pistils, the ovule-bearing structures, and stamens, the pollen-bearing structures. However, the two lateral spikelet flowers are normally sterile, lacking pistils, while the central spikelet maintains its fertility (Sakuma & Schnurbusch, 2020). In the barley mutant *six-rowed spike1* (*vrs1*), the lateral spikelets develop pistils that can be fertilized and develop into seeds. Natural mutations in *vrs1* were the source of the productive six-rowed barley (Komatsuda et al., 2007). Interestingly, an allele of *VRS1* that increases repression of pistil development in lateral spikelets can also increase yield by increasing the size of the mature grain in the central spikelet (Sakuma et al., 2017). Mutants of the *VRS1* homolog *GRAIN NUMBER INCREASE1 (GNI1)* also increased yield in wheat (*Triticum aestivum*) by increasing the number of fertile flowers per spikelet (Sakuma et al., 2019). Similarly, mutant alleles of the co-orthologs of *VRS1* increase kernel row number in maize (Kelliher et al., 2019). Thus, *VRS1* and similar genes regulate the growth of reproductive structures, in turn influencing yield.

Class I homeodomain-leucine zipper (HD-ZIP) transcription factors, encoded for by genes like *VRS1*, often regulate development via growth suppression (Akagi et al., 2014; Andres et al., 2017; González-Grandío et al., 2017; Kelliher et al., 2019; Komatsuda et al., 2007; Sakuma et al., 2019; Vlad et al., 2014; Vuolo et al., 2016; Whipple et al., 2011). Indeed, a *VRS1* homolog, *GRASSY TILLERS1* (*GT1*), suppresses growth of lateral branches in maize (Whipple et al., 2011). *gt1* plants develop tillers, have more ear-bearing branches, and sometimes develop pistils in the typically staminate tassel flowers (Whipple et al., 2011). In double mutants between *gt1* and the trehalose–phosphate phosphatase (TPP) gene *ramosa3* (*ra3*), pistil development in tassels is no longer suppressed, with receptive silks growing out of tassel flowers (Klein et al., 2022). In rice (*Oryza sativa*), an ortholog of *GT1* also represses lateral tillering when targeted by RNAi (V. Kumar et al., 2021). In arabidopsis (*Arabidopsis thaliana*), a triad of genes co-orthologous with *GT1* and *VRS1* (*HB21, HB40*, and *HB53*) similarly suppress lateral vegetative growth (González-Grandío et al., 2017). In persimmon (*Diospyros lotus*), the *GT1/VRS1* homologs *MeGI* and *OGI* jointly regulate sex determination through suppression of floral organs (Akagi et al., 2014). The genes *LATE MERISTEM IDENTITY1* (*LMI1*) and *REDUCED COMPLEXITY* (*RCO*), in a clade sister to the orthologs of *GT1* and *VRS1*, suppress leaf blade outgrowth along leaf margins in arabidopsis, *Cardamine hirsuta*, and cotton (*Gossypium hirsutum*) (Andres et al., 2017; Vlad et al., 2014; Vuolo et al., 2016).

Together, these data suggest that these class I HD-ZIP genes maintain a conserved and shared function of growth suppression.

To test this hypothesis, we used CRISPR-Cas9 gene editing and mutant analysis to ask what conserved and joint roles *VRS1* and *GT1* homologs have in regulating growth repression in the grasses. We discovered that these two genes jointly regulate growth suppression in maize and brachypodium. In double mutants, both vegetative and reproductive growth repression decreased. Because these genes likely have conserved roles across flowering plants, they represent a promising pathway for further targeted improvement of a broad range of crops.

## Results

### GT1 *and* VRS1 *homologs have complex gene orthology*

We first examined the evolutionary history of the *GT1* and *VRS1* homologs in the grasses to identify orthologous groups (Fig. 1A, Suppl. Fig 1). A series of duplications has led to complex genetic orthology in this gene clade. For example, the barley genes *VRS1* and *HvHOX2* are co-orthologous to a single gene in brachypodium because of a duplication shared by wheat and barley that excludes brachypodium (Sakuma et al., 2010).

**Figure 1:**
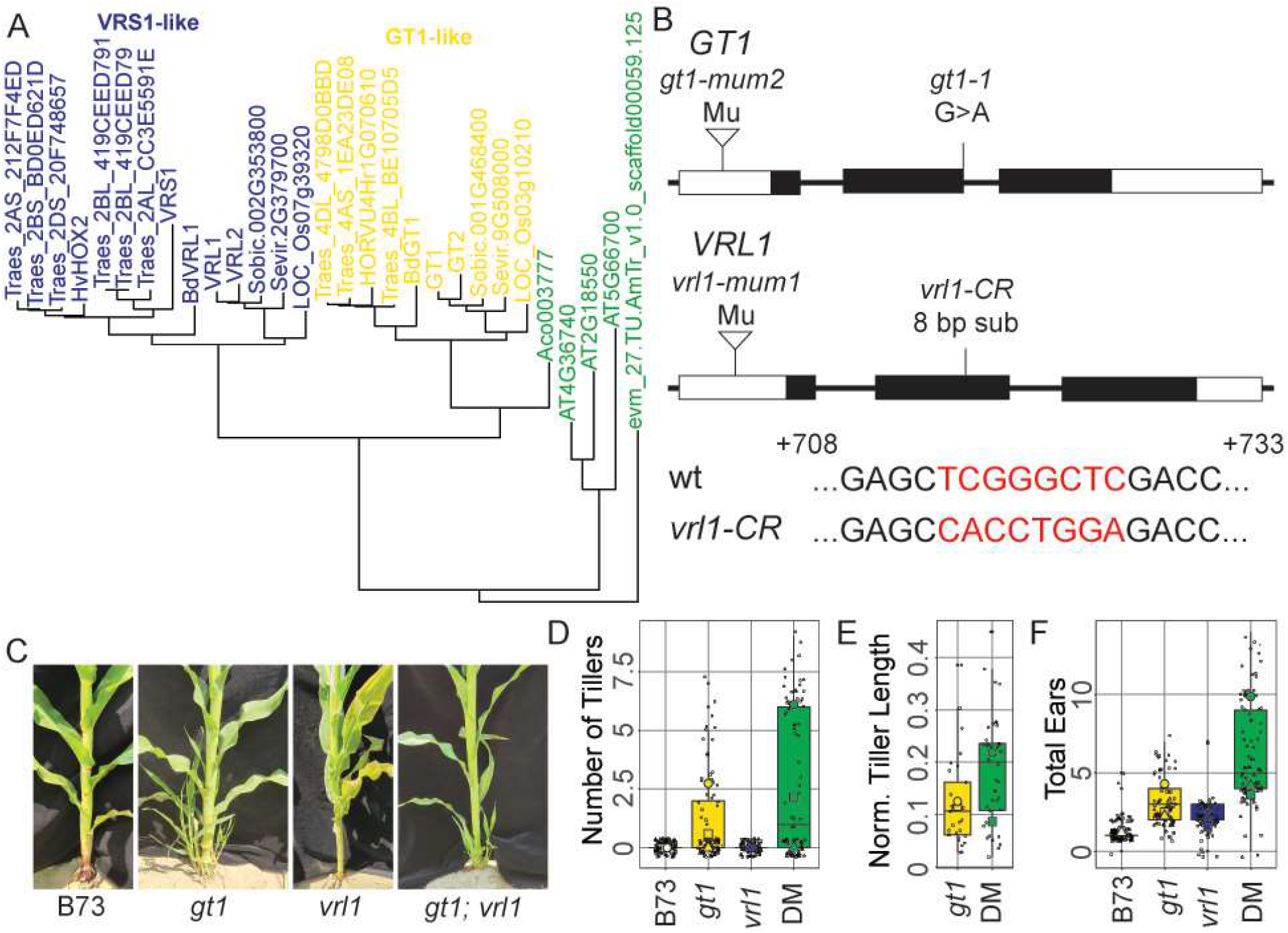
*VRL1* enhances *GT1* lateral growth repression. (A) Gene tree showing the evolutionary history of the *GT1* (yellow) and *VRS1* (blue) lineages of the class I HD-ZIPs. The black taxa are co-orthologous with *GT1* and *VRS1*. The black arrow points to the node at which the *VRS1* and *HvHOX2* lineages diverged. The gray arrows point to the nodes at which the maize genes *GT1/GT2* and *VRL1/VRL2* duplicated. Bootstrap support values are available in Supplementary Figure 1. (B) Mutant lesions used in this study. For *GT1*, a Mu-insertion line and the previously described *gt1-1* allele were used. For *VRL1*, a Mu-insertion line and a CRISPR mutant were used. An alignment of the wild-type *VRL1* allele and the *vrl1-CR* allele is provided. (C) Tillering in maize plants, left to right: wild-type, *gt1, vrl1, gt1; vrl1*. (D) Quantification of the number of tillers per plant. (E) Quantification of the height-normalized tiller length per plant. (F) Quantification of the number of ears produced per plant. In D-F, square = 2019 field, circle = 2020 field, triangle = 2021 field. Large shapes are year means, small points are individual plants.

Similarly, because of independent duplications in a lineage leading to maize (Swigonová et al., 2004), two maize genes are separately co-orthologous with *VRS1* and *HvHOX2*. Due to this complexity, we refer to the maize *VRS1* and *HvHOX2* co-orthologs as *VRS1-LIKE1* (*VRL1*) and *VRL2* and the brachypodium ortholog as *BdVRL1*.

### GT1 *and* VRL1 *exhibit joint roles in growth repression in ear flowers*

To test the roles of *GT1* and *VRL1* in the grasses, we examined the phenotypes in maize *Mutator* transposon insertion lines and in mutants generated via CRISPR-Cas9 genome editing (Fig 1B). *gt1* maize mutants have more tillers and ears compared to wildtype B73 (Fig. 1C,D,F; Whipple et al., 2011). In contrast, *vrl1* single mutants had more ears (but not more tillers) than wildtype B73 plants (Fig. 1C,D,F). To dissect the genetic interaction between *GT1* and *VRL1*, we made *gt1; vrl1* double mutants, which had even more ears and tillers than single mutants (Fig. 1C,D,F). In addition, *gt1; vrl1* tillers were longer than those of *gt1* (Fig. 1E). Together, these data suggest *GT1 and VRL1* are acting jointly to repress lateral branching in maize.

Because of the known floral suppression regulated by *GT1* and *VRS1* homologs in grasses (Klein et al., 2022; Komatsuda et al., 2007; Sakuma et al., 2019; Whipple et al., 2011), we also investigated changes in maize flowers. Maize has two inflorescence types: the staminate tassel at the apex of the plant, and the pistillate ears in the axils of upper leaves. In wildtype maize ears, spikelets initiate a pair of developing flowers. However, the lower flower aborts, in part because of pistil suppression (Fig 2A,B). This is also the case in *gt1* and *vrl1* single mutants (Fig 2C,D). Surprisingly, in *gt1; vrl1* double mutants, the pistil of the lower flower did not abort (Fig. 2E,F). Although this led to misrowing in *gt1; vrl1* ears, there was not a significant change in kernel row number (Suppl Fig. 2).

**Figure 2:**
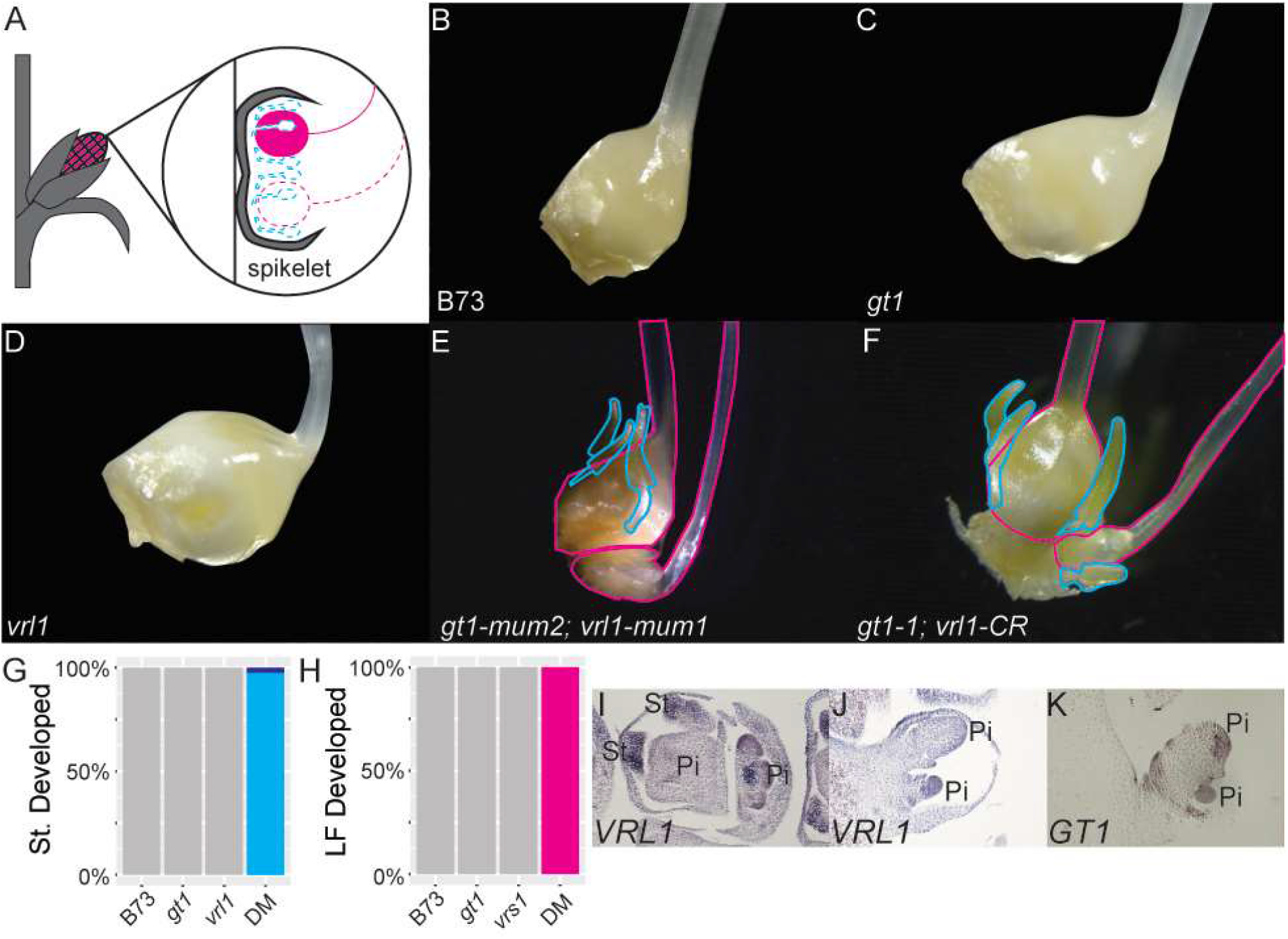
*VRL1* and *GT1* regulate repression of reproductive structures in the ear spikelets. (A) Diagram of maize ear spikelet development. In normal ears, the lower flower pistil (pink) and the stamens (cyan) of both flowers are repressed (dotted-line) while the upper flower pistil develops (solid). (B-F) Images of developing maize ear spikelets with palea, lemma, and lodicules removed. B, inbred line B73. C, *gt1*. D, *vrl1*. E, *gt1-mum2; vrl1-mum1*. F, *gt1-1; vrl1-CR*. G. Quantification of the number of ear spikelets with stamen-like structures. No wildtype, *gt1*, or *vrl1* plants developed stamen-like structures. All *gt1; vrl1* plants showed stamen-like structures in both flowers (cyan) except one, which showed stamen-like structures in the upper flower only (dark blue). H. Quantification of the number of ear spikelets with developed lower flower pistils. No wildtype, *gt1*, or *vrl1* plants developed lower flower pistils. All *gt1; vrl1* plants developed lower flower pistils. I-K. *in situ* hybridizations of *VRL1* (I, J) and *GT1* (K) in developing ear inflorescences. Pi = pistil, St = stamen.

Interestingly, the strong *gt1; vrl1* reproductive phenotype seems to be limited to ears rather than tassels. When initially examined, *gt1-mum2* and *gt1-mum2; vrl1-mum1* mutants bore long silks in the tassel flowers (Suppl. Fig. 3). However, upon backcrossing to B73, the strength of this phenotype decreased to that described previously for *gt1* (Klein et al., 2022; Whipple et al., 2011). This suggests that the silky tassel phenotypes of double mutants are subject to the regulation of a modifier in the initial background that is not present in the B73 background (Suppl Fig 3).

While wildtype and single mutants did not develop stamens in ear flowers, *gt1; vrl1* double mutants had stamen-like structures in both the upper and lower flower (Fig. 2E-H). These structures appeared similar to stamens, bearing structures that resembled anthers and filaments, but did not fully develop or produce pollen. To assess the genetic basis of these “stamens”, we examined mutants of more members of the *GT1/VRS1* gene clade in maize. We used CRISPR-Cas9 gene editing to generate mutant alleles in the *GT1* and *VRL1* homologs *GT2* and *VRL2* (Fig 1A). We found that plants fixed at *gt2-CR* and *vrl1-CR* but segregating for *vrl2-CR* also displayed derepression of stamen-like structures (Suppl Fig. 4A,C). However, no stamens were observed in *gt1-1; vrl1-CR*/+ plants, although there was a slight dosage effect leading to partially developed lower flower pistils in a portion of the spikelets (Suppl Fig 4B-D). In addition, *GT1* and *VRL1* are expressed in developing ear stamens and pistils (Fig. 2I-K). These data suggest that the joint action of *GT1* and *VRS1* homologs is required to suppress pistils and stamens in maize ear flowers, but that additional genetic components play a role in stamen suppression.

### *Single meristem transcriptome sequencing shows that* GT1 *and* VRL1 *regulate expression of catabolic processes*

We further asked which genes *GT1* and *VRL1* regulate. We performed RNA-seq of maize ear inflorescences in wildtype, *gt1, vrl1*, and *gt1; vrl1* individuals from a segregating population, and determined which genes were differentially expressed (Suppl. Table 3). A total of 219 genes were differentially expressed between wildtype and all the genotypes (Suppl. Table 4). Of these genes, 39 differentially expressed (DE) genes were shared among comparisons with all three genotypes, and 5, 46, and 111 were unique to each of the individual comparisons of wildtype with *gt1, vrl1*, and *gt1; vrl1*, respectively (Suppl. Fig. 5A).

Since *gt1; vrl1* double mutants have a strong ear phenotype, we further investigated the genes that were uniquely differentially expressed in ears (Suppl Fig 5B). A large number of transcription factors, including C2H2, bZIP, homeodomain, MYB, and WRKY transcription factors, were differentially expressed, as were receptors, sugar transporters associated with programmed cell death, and genes associated with floral development (Suppl Fig 5B; Suppl Table 4). Gene Ontology (GO) enrichment analysis of the Biological Processes category revealed that those genes that were up-regulated in *gt1; vrl1* double mutant versus the wildtype showed enrichment for positive regulation of various biosynthetic processes (Suppl Table 5), while those genes that were down-regulated in *gt1; vrl1* versus the wildtype were enriched for transmembrane transport and various catabolic processes (Suppl Table 5). These individual genes and GO categories align with our expectations that, with the failure to repress development of the lower flower in *gt1; vrl1* plants, there would be reduced expression of genes associated with breaking down and reusing the cellular components from the normally repressed structure.

Floral organs are also derepressed in *gt1; ra3* double mutant tassels (Klein et al., 2022). We compared the gene expression changes accompanying the derepression of pistils in tassel inflorescences in *gt1; ra3* maize plants (Klein et al., 2022) with the gene expression changes observed here.

Only three genes were shared between the wildtype versus *gt1; vrl1* DE genes and the pre-pistil suppression stage wild-type versus *gt1; ra3* DE genes (Klein et al., 2022). This overlap increased to eight genes when compared with the mid-pistil suppression stage (Suppl. Fig. 5C). These results suggest that the genes regulating floral organ repression in ear flowers differ from those regulating floral organ repression in tassel flowers.

Because of floral organ repression during normal development, lower and upper ear flowers have distinct gene expression patterns in maize (Yang et al., 2022). Since the *gt1; vrl1* phenotype effectively turns the lower flower of the ear into an upper flower of the ear, the set of genes that were DE between the lower flower and upper flower in Yang et al. 2022 might mirror the DE genes between wildtype and *gt1; vrl1*. However, we found a low overlap of only three DE genes (Suppl. Fig. 5C). This discrepancy is most likely due to biology and scale: the upper and lower flower data were collected prior to stamen initiation and targeted the floral meristem (Yang et al., 2022). Here, we collected whole inflorescence meristems at the stage when floral abortion would be taking place in the lower flower. These data suggest that the genes regulated by *gt1* and *vrl1* to regulate floral abortion are not part of floral meristem development at an earlier stage.

### GT1 *and* VRS1 *homologs maintain growth repression functions over deep time*

Maize is in a separate subfamily of grasses from barley and wheat, separated by ∼59 million years of evolution (S. Kumar et al., 2022). To further test our hypothesis that *GT1* and *VRS1* homologs have conserved roles in regulating growth repression, we made CRISPR-Cas9 knockouts of their homologs in the tractable model system brachypodium (Fig. 3A,B). *bdgt1* and *bdgt1; bdvrl1* both had more tillers than wildtype Bd21-3 (Fig. 3C-H). Spikelet number and flower number were both higher in *bdgt1; bdvrl1* double mutants than in either single mutant or wildtype (Fig. 3I,J). Despite the higher number of spikelets and flowers, neither seed number nor total seed weight per plant was significantly higher in any mutants compared to wildtype (Fig. 3K,L). This suggests that *BdGT1* and Bd*VRL1* play a role in repressing floral initiation and development, but there are additional genetic or environmental components that impact floral maturation into seed.

**Figure 3:**
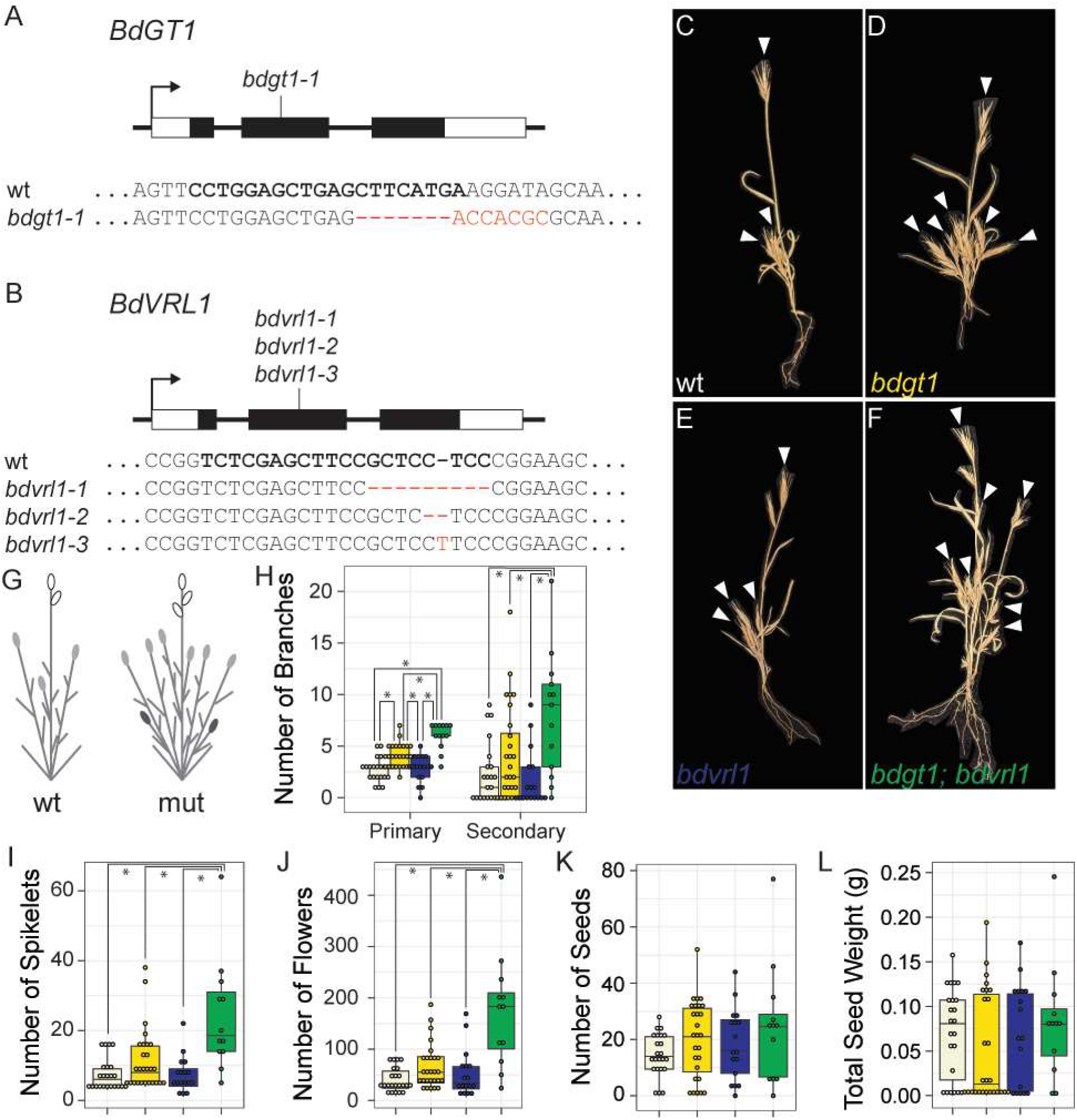
The growth repression role of *VRS1* and *GT1* homologs is conserved in brachypodium. (A, B) *bdgt1* and *bdvrl1* CRISPR lesions generated in this study. In the alignment, the bold text shows the site of the spacer, and the red text shows the changes from the wild-type sequence in each mutant allele. (C-F) The branching patterns of wild-type, *bdgt1, bdvrl1*, and *bdgt1; bdvrl1* brachypodium. White arrows point to spikelets on branch tips. (G) A diagram of the wild-type vs mutant brachypodium. White spikelets (ovals) are on the main axis, light gray spikelets are on the primary branches, and dark spikelets are on the secondary branches. (H) Quantification of the number of primary and secondary branches per plant. (I) Quantification of the number of spikelets per plant. (J) Quantification of the number of flowers per plant. (K) Quantification of the number of seeds per plant. (L) Quantification of the total seed weight per plant. Despite the increase in the number of flowers and seeds in the mutants, the total seed weight was the same on average. For H-L, stars denote *p*-value < 0.05.

Because of the repeated evolution of roles in growth repression in this gene lineage, we hypothesized that this lineage of HD-ZIPs could be recruited to repress growth in new contexts. To assess whether *GT1* and *VRS1* homologs were sufficient to repress growth in heterologous developmental contexts, we transgenically expressed maize *GT1* under the B-class gene *APETALA3* (*AP3*) promoter in arabidopsis (Jack et al., 1992; Lamb et al., 2002) (Fig. 4A). *AP3* specifies petal and stamen identity in arabidopsis flowers. If *GT1* could act as a growth repressor in a new context, we expected that this heterologous expression of *GT1* would reduce the size of petals and stamens (Fig 4A). Indeed, both petal area and stamen length were significantly reduced in the *AP3::GT1* plants compared to the control *AP3::YFP* plants (Fig. 4B,C). This demonstrates that, even across 160 million years of evolutionary time, *GT1* may be recruited for initiating growth repression (S. Kumar et al., 2022).

**Figure 4:**
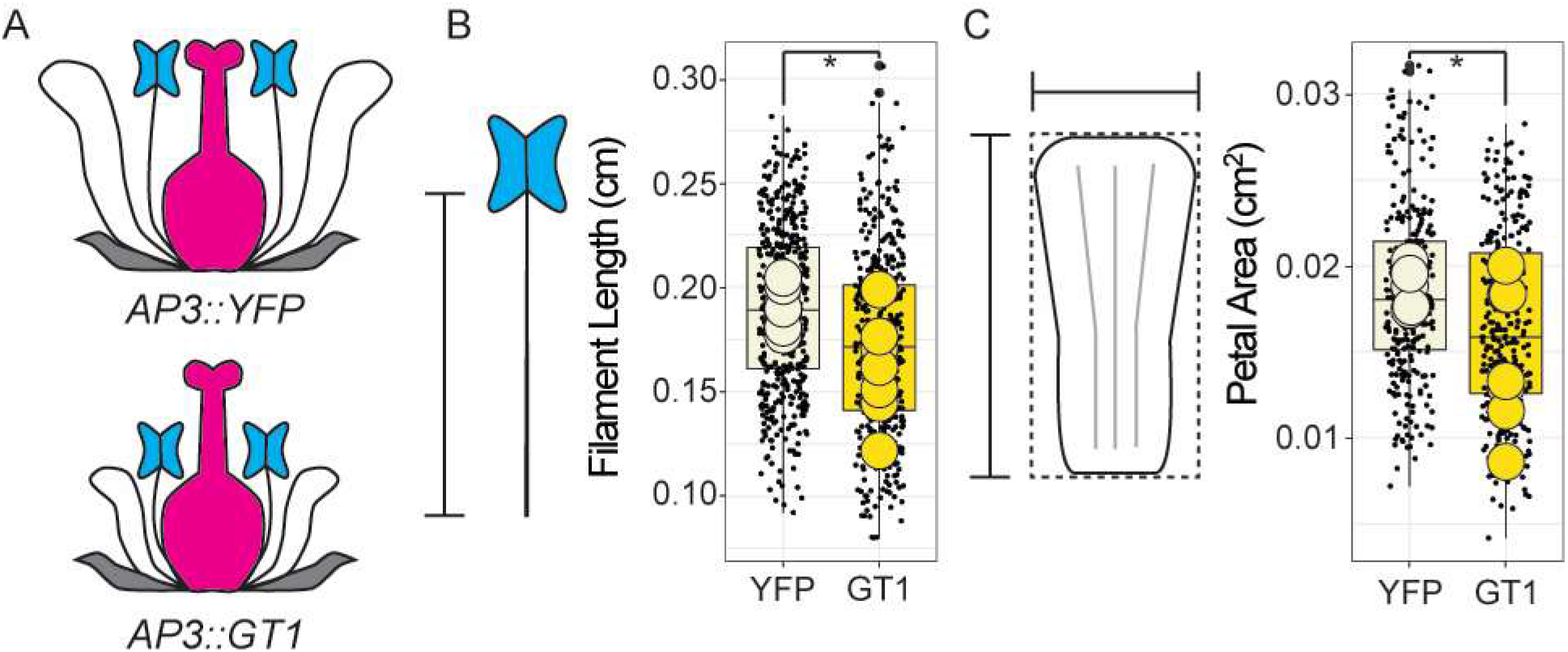
The conserved growth repression role of GT1 is maintained in new contexts. (A) Floral diagram showing the expected phenotype of transgenic arabidopsis flowers with either *AP3::YFP* or *AP3::GT1*. With *AP3::YFP*, we expect normal growth. With *AP3::GT1*, we expect that tissues that express *AP3* normally (petals and stamens) will be smaller due to the role of *GT1* in growth repression. (B) Quantification of the length of filaments in transgenic arabidopsis. The filament length is shorter in *AP3::GT1* compared to *AP3::YFP* (p-value = 8.943e-09). (C) Quantification of the petal area in transgenic Arabidopsis. The petal area is smaller in *AP3::GT1* than in *AP3::YFP* (p-value = 4.903e-06).

## Discussion

Here, we show that *GT1-*like and *VRS1-*like genes have undergone evolutionary divergence for *ca*. 80 My, but that they both perform roles in growth suppression in the grasses (Fig 1A, S. Kumar et al., 2022). Although maize *vrl1* plants do not exhibit obvious phenotypes, maize *gt1; vrl1* double mutants develop more ears and more floral organs in the ear spikelets than wildtype or *gt1* mutants (Fig 1F, Fig 2). Single meristem transcriptomics showed that *gt1; vrl1* plants display lower expression of genes involved in catabolism (Suppl Table 5). Gene-edited brachypodium had increased numbers of branches, spikelets, and flowers in *bdgt1; bdvrl1* plants, confirming these genes’ role in growth repression (Fig 3H,I,J). Thus, *GT1*-like and *VRS1*-like genes maintain a conserved function to jointly regulate growth suppression of developmental traits in the grasses. Further, ectopically expressed *GT1* suppressed growth of arabidopsis floral organs, demonstrating that these genes can regulate growth suppression even across much deeper timescales (Fig 4B,C).

Flower number and fertility are key yield-related traits regulated by *VRS1* in barley and its homolog *GNI1* in wheat (Komatsuda et al., 2007; Sakuma et al., 2019). Here, in line with that data, the double mutants in maize and brachypodium both showed an increase in flower number (Fig. 2E, F, Suppl. Fig. 2A, Fig. 3J). Yet, in our study, this increase in flowers did not necessarily lead to a change in yield. Kernel row number in maize was unchanged in *gt1; vrl1* mutants (Suppl. Fig 2B), and *bdgt1; bdvrl1* mutants did not produce more seeds (Fig. 3K,L). While *GT1* and *VRS1* homologs may regulate the number of flowers that develop, additional genetic components may contribute to the maturation of those flowers into seeds. In barley, the *CCT MOTIF FAMILY4* (*CMF4*) gene modulates spikelet survival and maturation through the vascular-associated circadian clock and organ greening (Huang et al., 2023). To maximize the yield-related effect of developing or selecting *GT1* and *VRS1* homolog alleles that permit more floral development, concurrent selection of floral maturation genes like *CMF4* may need to occur.

### *The conserved role of* GT1*-like and* VRS1*-like genes in growth repression facilitates their recruitment for growth repression in new contexts*

*GT1-*like and *VRS1-*like genes have conserved roles in growth repression across species, but that repression can occur in different contexts. Mutations in these genes in maize and brachypodium increased vegetative branching relative to wildtype (Fig. 1, 3). This conservation is reinforced by the shared role of the three arabidopsis co-orthologs in lateral branching (González-Grandío et al., 2017). However, in the grasses, including maize and brachypodium, *GT1* and *VRS1* genes diverged and took on new roles in regulating floral development (Fig. 2, 3; Klein et al., 2022; Whipple et al., 2011). This has also happened in barley, where *VRS1* controls the development of the carpels in lateral spikelets (Komatsuda et al., 2007) and in wheat, where *GNI1* regulates the number of fertile flowers within a spikelet (Sakuma et al., 2019). Outside of the grasses, *GT1/VRS1* homologs *MeGI* and *OGI* have developed an antagonistic relationship to control sexual differentiation via growth repression in persimmon (Akagi et al., 2014). A similar mechanism of growth repression control has also been proposed for *VRS1* and its barley-specific ortholog *HvHOX2* (Sakuma et al., 2010; Thirulogachandar et al., 2021). These examples of *GT1*-like and *VRS1*-like genes show how these genes have been repeatedly recruited to work in new contexts yet retain their capacity to repress growth across flowering plants.

These repeated and wide-spread examples of the *GT1*-like and *VRS1*-like genes changing or expanding their roles to regulate growth repression in new contexts demonstrates their propensity for recruitment into new developmental contexts. These data together suggest that this group of class I HD-ZIPs may be “genetic hotspots” that are involved frequently in morphological change (Martin & Orgogozo, 2013; Stern & Orgogozo, 2009). Why would these genes be predisposed to regulating morphological change? We propose that *GT1* and *VRS1* occupy a conserved position within the genetic network that allows them to regulate morphological characters without a high level of pleiotropy, and thus they may be more likely to gain and retain new roles within development (Martin & Orgogozo, 2013; Stern & Orgogozo, 2009; Uller et al., 2018; Wang et al., 2022). The capacity of these genes to repress growth across diverse species also supports this idea (Fig. 1, 2, 3, 4). Even across 160 million years of evolutionary time, maize *GT1* can suppress growth in arabidopsis floral tissues, showing that expression of *GT1* is sufficient to repress growth in new contexts (S. Kumar et al., 2022). Even *LMI* and *RCO* genes from the sister clade to *GT1*- and *VRS1*-like genes show evidence of regulating growth repression (Vlad et al., 2014; Vuolo et al., 2016). Given the various roles for which *GT1* and *VRS1* genes have been recruited, it seems likely that these genes are hotspots predisposed to roles in growth repression regulation in new contexts (Akagi et al., 2014; Klein et al., 2022; Komatsuda et al., 2007; V. Kumar et al., 2021; Sakuma et al., 2019; Whipple et al., 2011). To evaluate this hypothesis, further work confirming that regulators of *GT1/VRS1* homologs vary while downstream function remains relatively constant across diverse species would be necessary.

### *Following evolution’s lead to improve crop engineering and* de novo *domestication*

*De novo* domestication involves the improvement of wild crop relatives or partially domesticated crop species using gene editing (M. E. Bartlett et al., 2022). Progress has been made in a handful of species, but the full process of domestication is complex (Kwon et al., 2020; Lacchini et al., 2020; Lemmon et al., 2018; H. Yu et al., 2021; Zsögön et al., 2018). To accelerate and improve the efficiency of *de novo* domestication, criteria for selecting the best targets for genome editing will need to be established.

One criterion that could improve *de novo* domestication is targeting genes with conserved functions. Genes with conserved roles across species are more likely to regulate the same trait in any given species. For example, the well-conserved meristem maintenance pathway is regulated by the *CLAVATA1* (*CLV1*) and *CLAVATA3* (*CLV3*) genes in arabidopsis, and orthologous genes in other species have been found to regulate this pathway in a similar fashion (Fletcher, 2018). In maize and tomato, when a series of gene edited alleles were developed that modified the upstream regulatory region of *CLV3* orthologs, meristem size increased, resulting in kernel row number and fruit size increase, respectively (Liu et al., 2021; Rodríguez-Leal et al., 2017). Similarly, in tomato, *SELF PRUNING* (*SP*) and *SELF PRUNING 5G* (*SP5G*) belong to a gene family with conserved roles in flowering repression across angiosperms (Pnueli et al., 1998), and these genes were successfully modified to increase determinacy and compactness and decrease time to flowering (Kwon et al., 2020). Thus, we should prioritize genes with well-conserved roles that regulate our specific trait of interest. The conservation of function in *GT1*-like and *VRS1*-like genes as well as their importance in domestication support their position as priority *de novo* domestication targets.

A second criterion for targeting a gene may be the likelihood of generating a beneficial phenotype in the resulting gene edited plant. While genes that are positive regulators may require increase of expression or a gain-of-function allele to develop a beneficial trait, genes that are negative regulators may permit positive traits via reduction in expression or loss-of-function. Thus, editing these genes is more likely to lead to an agronomically beneficial change, since loss of function alleles are easier to generate. This is likely one of the reasons for the success in improving traits via *CLV3* ortholog gene editing: reduction in *CLV3* ortholog expression led to an increase in meristem size, resulting in increased cob and fruit sizes (Liu et al., 2021; Rodríguez-Leal et al., 2017). Similarly, *GT1*-like and *VRS1*-like genes are negative regulators of growth. Mutants in these genes lead to increased growth that translates to an increase in the number of flowers (Fig 2H, Fig 3J). This increase in fertility makes them strong candidate targets for de novo domestication.

Another consideration when targeting genes for de novo domestication is the likely strength of alleles. Strong alleles may lead to more extreme phenotypes, but the development of the plant may be disrupted when incorporating that change.

In maize, mutations in several genes of the CLV pathway lead to strong fasciation phenotypes that would diminish their use in agriculture (Bommert et al., 2005; Je et al., 2016, 2018; Liu et al., 2021). Weak alleles may be more easily incorporated into plant development without drastic alterations. A series of weak alleles that modify the upstream elements of the maize *CLV3* orthologs displayed a spectrum of meristem size change, resulting in ears with increased kernel row number without malformation (Liu et al., 2021). Similarly, a weak allele of another CLV pathway gene led to an increase in yield in elite maize lines (Trung et al., 2020).

While the alleles of *GT1* and *VRS1* homologs examined here might be considered strong alleles, the presence of paralogs, gene duplicates within the same species, complicates the strength of mutant alleles. In tomato, the *CLV3* homolog has undergone duplication, and one copy plays a primary role in maintaining meristem size. However, knockouts of this gene led to paralogous compensation, whereby the second copy becomes more highly expressed when the first copy is knocked out (Rodriguez-Leal et al., 2019). A survey of these genes in other Solanaceae species showed that these species have variably lost one copy or retained both. Yet, even in cases where both copies have been retained, the patterns of paralog compensation are not always the same (Kwon et al., 2022). Similarly, the *GT1/VRS1-like* genes have a complicated history with duplication (Fig 1A), and may compensate for each other.

While paralogy and compensation may be considered a hindrance, it can also be a tool for creating even greater nuance in phenotypic changes. A combination of allele strengths in multiple *GT1*-like and *VRS1*-like genes might develop more moderate, but still beneficial, phenotypes and provide a more granular understanding of their distinct roles in vegetative and reproductive development. Dissecting their regulation via gene editing could provide a clearer view for how to use these genes for crop engineering and *de novo* domestication (M. Bartlett, 2019; M. E. Bartlett et al., 2022; Eshed & Lippman, 2019). This information can then be translated to tuning gene expression and function for crop improvement.

## Materials and Methods

### Gene tree inference

Protein sequences of GT1, VRS1 and known homologs were used as a BLASTP query against a peptide databases of 30 plant genomes available on Phytozome plus the *D. lotus* and *C. hirsuta* genome websites with an e-value cut-off of 10e-30 (Suppl Table 2). The peptide sequences that met the cut-off were aligned using MAFFT (Katoh et al., 2002). The alignment was filtered with noisy (Dress et al., 2008). A model selection test was performed and a tree was constructed in IQ-TREE 2 (Minh et al., 2020) and visualized using the R package ggtree (G. Yu, 2020). The outgroups for the tree were three HD-ZIP genes from classes II, III, and IV.

### Maize plant materials and growth conditions

Maize mutants with Mutator transposon insertions in the 5’ UTRs of *gt1* (Zm00001d028129) and one of the co-orthologous maize *vrs1* (Zm00001d021934) loci, here called *vrs1-like1 (vrl1)*, were identified via a reverse genetics screen as in (Bensen et al., 1995). These alleles were designated *gt1-mum2* and *vrl1-mum1*. Genotyping of *gt1-mum2* and *vrl1-mum1* alleles was performed via triplex PCR with a MuTIR primer and two gene-specific primers (Suppl Table 1).

Spacers targeting the maize homologs of *GT1* and *VRS1* were designed and assembled into pENTR and Gateway cloned into the Cas9-containing plasmid pMCG1005-Cas9 (Char et al., 2017). This construct was transformed into *Agrobacterium tumefaciens* strain EHA101. Plant transformation was performed at the Iowa State University Plant Transformation Facility (Ames, IA) into the HiII background. Regenerated plantlets were screened for herbicide resistance to detect successful transformants. CRISPR-Cas9 edited mutants of *vrl1-CR, vrl2*-CR (Zm00001d006687), and *gt2*-CR (Zm00001d048172) were genotyped via PCR and Sanger sequencing. Spacer sequences and primers are available in Suppl. Table 1.

Plants for vegetative and reproductive phenotyping and *in situ* hybridization were grown at the University of Massachusetts Amherst Crop and Animal Research and Education Farm in South Deerfield, MA (∼42°29′N, 72°35′W). Plants for RNA-seq were grown at the College of Natural Sciences and Education Greenhouse on the UMass Amherst campus under long day conditions (16 hours light, 8 hours dark) at 28°C.

### Maize phenotyping

We measured phenotypes in maize inbred line B73 and in *gt1-mum2, vrl1-mum1*, and *gt1-mum2; vrl1-mum1* plants in their native background. Plants for phenotyping were grown in four blocks for three years (2018, 2019, 2020). For each plant in each row, tiller number, tiller length, plant height, ears at each node, silks in tassels, and derepressed lower flowers were counted or measured. Tiller lengths were normalized by plant height for statistical comparisons. For counting derepressed lower flowers and floral organs, ten spikelets per plant, four plants per genotype, were examined under the dissecting microscope. To establish if tassel silks were a background effect, *gt1-mum2; vrl1-mum1* plants were backcrossed into the B73 inbred background four times, selfed, and phenotyped. Previously described *gt1-1* (Whipple et al., 2011) and CRISPR alleles *vrl1-CR* were used to confirm initially observed phenotypes. These alleles plus CRISPR alleles of genes *vrl2-CR* and *gt2-CR* were used to phenotype the presence of stamen-like structure in maize ears.

### CRISPR-Cas9 genome editing in brachypodium

Spacers targeting *BdGT1* (Bradi1g71280) and *BdVRL1* (Bradi1g23460) were designed using the software CRISPOR (Concordet & Haeussler, 2018). These spacers were synthesized and assembled into a guide RNA construct using the MoClo system (Engler et al., 2014; Weber et al., 2011). The resulting construct was Gateway cloned into pOsCas9_RC_of_L, a Cas9- and hygromycin resistance-containing expression vector (Miao et al., 2013; O’Connor et al., 2017). This construct was introduced to brachypodium Bd21-3 embryogenic callus via *Agrobacterium*-mediated transformation using strain AGL1 (Vogel & Hill, 2008). Calli were screened for successful transformation via hygromycin resistance, and were regenerated into plantlets. *bdgt1* and *bdvrl1* alleles were genotyped via PCR and Sanger sequencing. Spacer sequences and primers are available in Suppl. Table 1.

Brachypodium plants were grown in growth chambers either at the Morrill Greenhouse on the UMass Amherst campus or in the Bartlett lab. All plants were grown under long day conditions (20 hrs light, 4 hrs dark) at 24 °C.

### In situ hybridizations

Maize B73 ears were fixed, embedded, sectioned, and hybridized as described in (Jackson et al., 1994). Detection incubation was 24 to 48 hrs long. Hybridizations were performed using the *GT1* antisense digoxigenin-labeled RNA probe described by (Whipple et al., 2011). *VRL1* cDNA was amplified using primers in Suppl Table 1 and was cloned into pJET1.2/blunt (Thermofisher). Antisense digoxigenin-labeled RNA probes were synthesized using MEGAscript T7 Transcription Kit (Invitrogen).

### RNA-seq sampling and analysis

From a family segregating for *gt1-mum2* and *vrl1-mum1* and backcrossed into B73 inbred background four times, 56 maize ears were collected and frozen in liquid nitrogen. RNA was extracted from these tissues using Qiagen RNeasy Plant Mini Kit (Qiagen), treated with DNase I, and cleaned up with Monarch RNA Cleanup Kit (10 ug) (NEB). These cleaned RNA samples were sent to Novogene for poly-A tail enrichment mRNA library preparation and 150 bp paired end sequencing on Illumina NovaSeq (Novogene, Sacramento, CA, USA).

Sequenced RNA libraries were trimmed for quality using Trimmomatic and mapped to the *Zea mays* (maize) B73 version 5 genome using STAR 2.7.9a (Dobin et al., 2013; Hufford et al., 2021). Reads mapped to genes were counted using HTSeq (Anders et al., 2015). RUVSeq was used to normalize the read counts using upper quartile normalization and the expression of the 5000 least differentially expressed genes between the *gt1; vrl1* and wildtype ear samples (Risso et al., 2014). Differential expression analysis was performed using DESeq2 (Love et al., 2014). Gene ontology analysis was performed on differentially expressed gene sets using topGO (Alexa & Rahnenfuhrer, 2016).

### Arabidopsis constructs and transformation

The promoter of arabidopsis *APETALA3* (*AP3*) and the coding sequence of maize *GT1* were amplified and cloned into backbones using the MoClo System (Engler et al., 2014; Weber et al., 2011) (Suppl Table 1). These were Golden Gate cloned into kanamycin-resistant expression vector pICH86966 with the CaMV 35S terminator (pICH41414) (Engler et al., 2014; Weber et al., 2011). This construct was transformed into arabidopsis using the floral dip method of *Agrobacterium*-mediated transformation in strain GV3101, as in Zhang et al. (2006). Transformed arabidopsis were screened for kanamycin resistance. Resistant T0 generation plants were phenotyped for petal and stamen size in opened flowers.

## Data Availability

Raw sequencing data are available at the National Center for Biotechnology Information BioProjects (PRJNA######); Data sets underlying the figures are available in supplementary data sets. Code for analyses is available on GitHub (https://github.com/BartlettLab/GT1VRS1)

## Acknowledgements

This work was supported by an NSF CAREER award (IOS-1652380) to M. E. Bartlett, and a USDA NIFA AFRI Postdoctoral Fellowship (2019-67012-29654) to J. P. Gallagher.

## Supplementary Figures and Tables

**Supplementary Figure 1. GT1and VRS1 gene tree**. *GT1* homologs are colored in goldenrod, VRS1 homologs are colored in blue, and gene co-orthologous with *GT1* and *VRS1* are colored in green. The genes used in this study are marked with their gene symbol. Numbers at nodes are bootstrap percentages.

**Supplementary Figure 2. Misrowing and kernel row number in *gt1* and *vrl1* mutants**. A. *gt1-mum2; vrl1-mum1* shows misrowing of kernels due to derepression lower flowers in maize ears. B. Despite misrowing, kernel row number (KRN) does not vary among *gt1-mum2, vrl1-mum1*, or *gt1-mum2; vrl1-mum1*.

**Supplementary Figure 3. The tassel silks associated with *gt1-mum2* is due to a background effect**. (A-F) Whole tassel phenotype for B73, *gt1-1* in B73, *gt1-mum2, vrl1-mum1, gt1-mum2; vrl1-mum1*, and *gt1-mum2; vrl1-mum1* backcrossed into B73 four times (left to right). Only individuals homozygous for *gt1-mum2* bore silks in tassels. (G-L) Tassel floret phenotype for B73, *gt1-1* in B73, *gt1-mum2, vrl1-mum1, gt1-mum2; vrl1-mum1*, and *gt1-mum2; vrl1-mum1* backcrossed into B73 four times (left to right). B73 and *vrl1-mum1* do not show any developing tassel carpels, while *gt1-1* shows a very small tassel carpel (red box). *gt1-mum2* and *gt1-mum2; vrl1-mum1* both show a more fully developed tassel carpel. However, the backcrossed *gt1-mum2; vrl1-mum1* tassel floret does not show carpel development.

**Supplementary Figure 4. Stamen repression in ear florets involves multiple *GT1* and *VRS1* homologs**. A. Individuals fixed for *gt2* and *vrl1* produce stamen-like structures in ear flowers. B. *gt1; vrl1*/+ individuals do not produce stamen-like structures but do occasional derepression of small pistils. C. Quantification of stamen-like structures in ear flowers. D. Quantification of lower ear flower pistil development. The lighter pink color denotes counts of small pistils, as in B, second panel. E. *in situ* hybridizations of *VRS1* (panel 1, 2) and *GT1* (panel 3) in developing ear inflorescences. Pi = pistil, St = stamen.

**Supplementary Figure 5. Heatmap of RNA-seq data**. A. Venn diagram of genes differentially expressed in *gt1, vrl1*, and *gt1; vrl1* versus wildtype. B. Heatmap of the 85 genes significantly differentially expressed between wild-type and *gt1; vrl1* with |logFC| greater than 1. C. Number of overlapping genes between this study and those from previous studies.

**Supplementary Table 1. Primers and oligos used in this study. Supplementary Table 2. Genomes used for BLASTP search**.

**Supplementary Table 3. RNA-seq library sequencing statistics**

**Supplementary Table 4. Gene differentially expressed between wildtype and *gt1, vrl1*, and *gt1;vrl1***

**Supplementary Table 5. Gene Ontology analysis of significantly up- and down-regulated genes between wildtype and *gt1; vrl1***.

