## Supplementary figures and images for "Duplicate transcription factors *GT1* and *VRS1* regulate branching and fertile flower number in maize and *Brachypodium distachyon*"

### Supplemental Figure 1

**Supplementary Figure 1.** ***GT1*** **and** ***VRS1*** **gene tree**

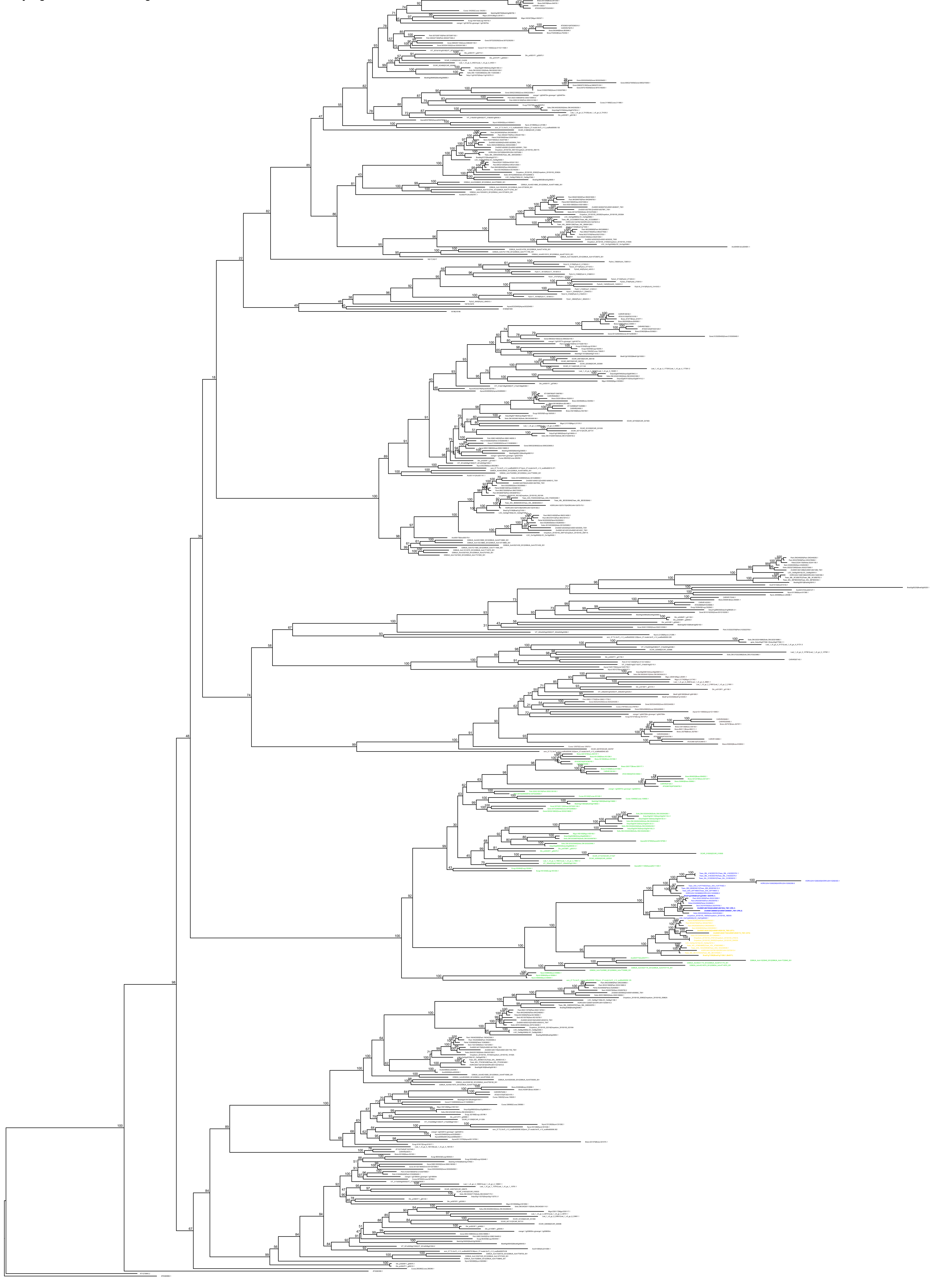
