## Supplemental Figure 2 for "Duplicate transcription factors *GT1* and *VRS1* regulate branching and fertile flower number in maize and *Brachypodium distachyon*"

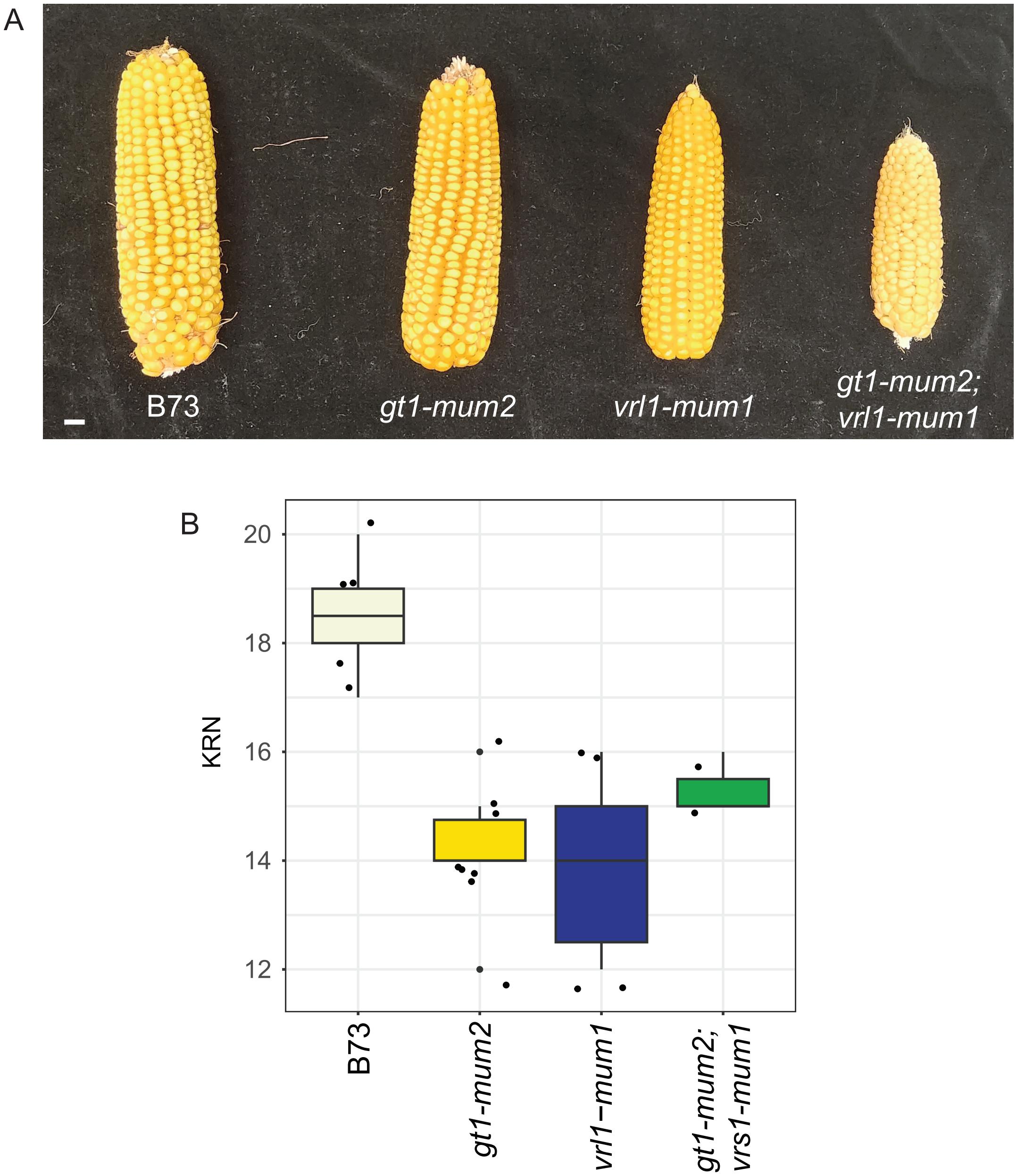


Supplementary Figure 2. Misrowing and kernel row number in gt1 and vrl1 mutants. A. *gt1-mum2; vrl1-mum1* shows misrowing of kernels due to derepression lower flowers in maize ears. B. Despite misrowing, kernel row number (KRN) does not vary among *gt1-mum2*, *vrl1-mum1*, or *gt1-mum2; vrl1-mum1.*
