## Supplemental Figure 3 for "Duplicate transcription factors *GT1* and *VRS1* regulate branching and fertile flower number in maize and *Brachypodium distachyon*"

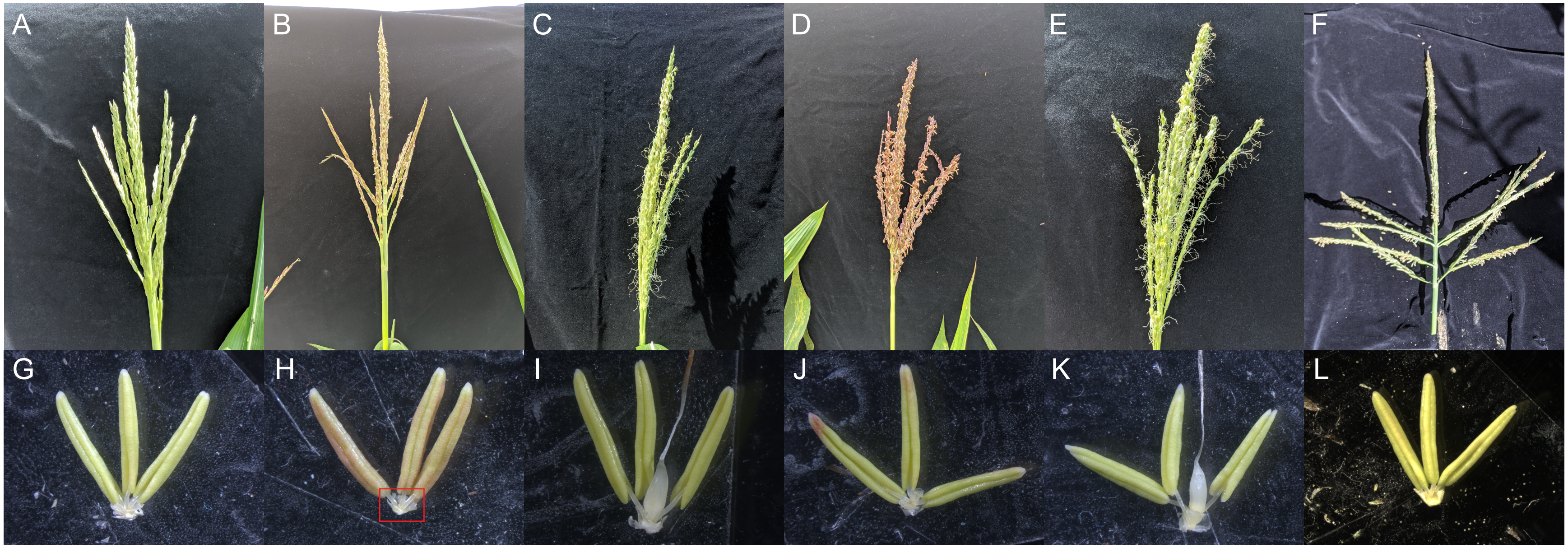


Supplementary Figure 3. The tassel silks associated with *gt1-mum2* is due to a background effect. (A-F) Whole tassel phenotype for B73, *gt1-1* in B73, *gt1-mum2*, *vrl1-mum1, gt1-mum2; vrl1-mum1,* and *gt1-mum2; vrl1-mum1* backcrossed into B73 four times (left to right). Only individuals homozygous for *gt1-mum2* bore silks in tassels. (G-L) Tassel floret phenotype for B73, *gt1-1* in B73, *gt1-mum2*, *vrl1-mum1*, *gt1-mum2; vrl1-mum1*, and *gt1-mum2; vrl1-mum1* backcrossed into B73 four times (left to right). B73 and *vrl1-mum1* do not show any developing tassel carpels, while *gt1-1* shows a very small tassel carpel (red box). *gt1-mum2* and *gt1-mum2; vrl1-mum1* both show a more fully developed tassel carpel. However, the backcrossed *gt1-mum2; vrl1-mum1* tassel floret does not show carpel development.
