## Supplemental Figure 4 for "Duplicate transcription factors *GT1* and *VRS1* regulate branching and fertile flower number in maize and *Brachypodium distachyon*"

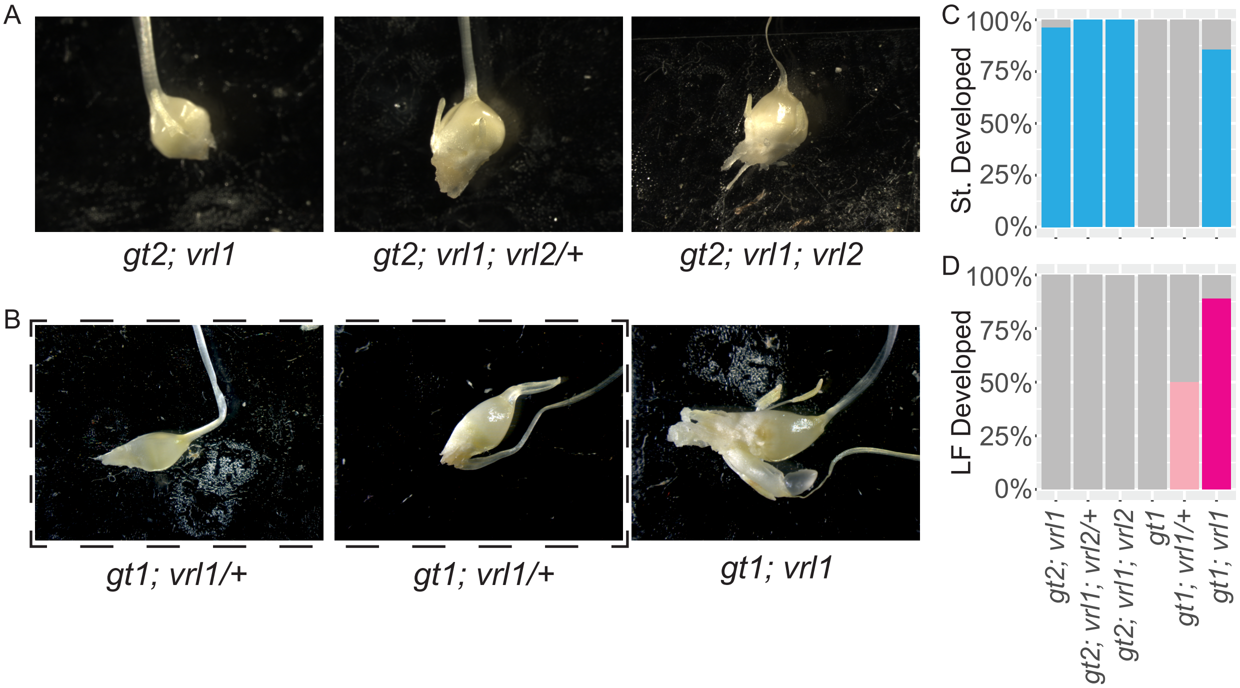


Supplementary Figure 4. Stamen repression in ear florets involves multiple *GT1* and *VRS1* homologs. A. Individuals fixed for *gt2* and *vrl1* produce stamen-like structures in ear flowers. B. *gt1; vrl1/+* individuals do not produce stamen-like structures but do occaisonal derepression of small pistils. C. Quantification of stamen-like structures in ear flowers. D. Quantification of lower ear flower pistil development. The lighter pink denotes counts of small pistils, as in B, second panel.
