## Supplemental Figure 5 for "Duplicate transcription factors *GT1* and *VRS1* regulate branching and fertile flower number in maize and *Brachypodium distachyon*"

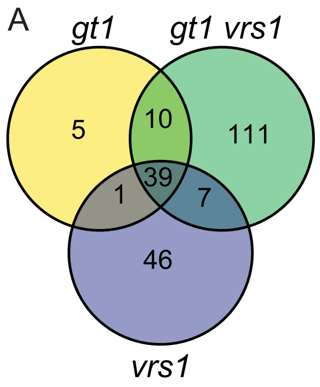


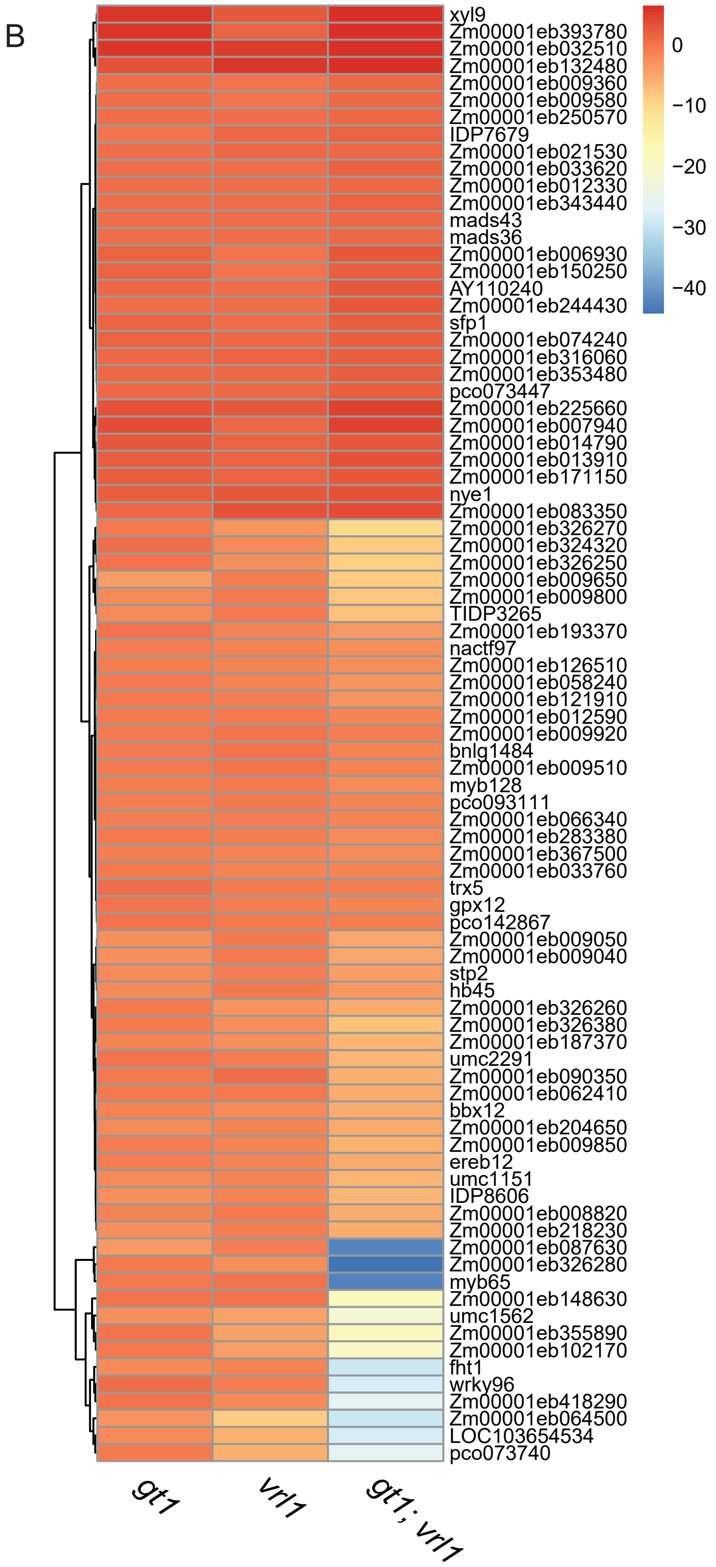


C

| Study | Up/Down |
|---|---|
| Yang *et al.* 2021 | 3/0 |
| Klein *et al.* 2022 | |
| pre-suppression | 2/1 |
| mid-suppression | 3/5 |

Supplementary Figure 5: A. Venn diagram of genes differentially expressed in *gt1*, *vrl1*, and *gt1; vrl1* versus wildtype. B. Heatmap of the 85 genes significantly differentially expressed between wild-type and *gt1; vrl1* with |logFC| greater than 1. C. Number of overlapping genes between this study and those from previous studies.
