## Supplemental Table 1 for "Duplicate transcription factors *GT1* and *VRS1* regulate branching and fertile flower number in maize and *Brachypodium distachyon*"

**Supplementary Table 1: Primers and oligos used in this study**

| **Genotyping primers** | | | |
| --- | --- | --- | --- |
| **Name** | **Sequence** | | **Source** |
| gt1-mum2-F | TCTCCGGCCGGCCTATAAAT | | This study |
| gt1-mum2-R | GCCAGTCAATCCTTCCTTCGTCTTGTAG | | This study |
| vrl1-mum1-F | TTTTACATCCATAGCCGGTCTGCCTCAGA | | This study |
| vrl1-mum1-R | GAAGCTCAGCTCCAGCATCTCTACCT | | This study |
| MuTIR | AGAGAAGCCAACGCCAWCGCCTCYATTTCGTC | | This study |
| GT1-CR_F | CTACCAGGTGGTGAGCAGAA | | This study |
| GT1-CR_R | ATACCACGCCGAGGACACTC | | This study |
| GT2-CR_1 | GATGTACTTATATGTGCTACCAGGTAGAC | | This study |
| GT2-CR_R | TCACTATTATGTGGTCGTGTCG | | This study |
| VRL1-CR_F | TGCTACCGGCTGCTAGGAA | | This study |
| VRL1-CR_R | TGCGGGCAGGGCAGTATTAC | | This study |
| VRL2-CR_F | CATTCTCTTTCTCACAGTTGTGT | | This study |
| VRL2-CR_R | CAGGCAAAGAGTGAGAGGT | | This study |
| gt1-1_F | AGGTGGCCGTCTGGTTCCAGAA | | Whipple et al. 2011 |
| gt1-1_R | TGGTGCGTCACCGTCGAGAAC | | Whipple et al. 2011 |
| **CRISPR spacers** | | | |
| **Name** | | **Sequence** | **Source** |
| GT1VRS1-1 | | GCTTGCTCTTGTGGCGGGCG | This study |
| GT1VRS1-2 | | GTGCACCTGGCCGCCGAGCT | This study |
| BdGT1_341forw | | TATCCAAGCTGCAAAGCCCA | This study |
| BdGT1_1083rev | | TGGCTAGCTAACCGCCAAAA | This study |
| BdVRL1_1037forw | | TCGAGGAGCTTGCTCTTGTG | This study |
| BdVRL1_1584rev | | TGGTGGTCCTAGCTCCTCTC | This study |
| **In situ probe primers** | | | |
| Name | Sequence | | Source |
| HDLZ-For | CCTAGTCCTAGTACAGGCTACAG | | Whipple et al. 2011 |
| HDLZ-Rev | CGGTCCATCCATCCATTAACACG | | Whipple et al. 2011 |
| VRL1_probe_F1 | AGGAGAGAGAAAGGCGTTGC | | This study |
| VRL1_probe_R1 | TAATCTGCCGACGACGATGG | | This study |
