## Supplemental Table 2 for "Duplicate transcription factors *GT1* and *VRS1* regulate branching and fertile flower number in maize and *Brachypodium distachyon*"

**Supplementary Table 2: Genomes used for BLASTP search**

| **Species Name** | **Genome Version** | **Source** | **Website** | **Reference** |
| --- | --- | --- | --- | --- |
| *Amborella trichopoda* | v1.0 | Phytozome 13 | https://phytozome-next.jgi.doe.gov/ | <https://doi.org/10.1126/science.1241089> |
| *Ananas comosus* | v3 | Phytozome 13 | https://phytozome-next.jgi.doe.gov/ | <https://doi.org/10.1038/ng.3435> |
| *Aquilegia coerulea* | v3.1 | Phytozome 13 | https://phytozome-next.jgi.doe.gov/ | <https://doi.org/10.7554/elife.36426> |
| *Arabidopsis thaliana* | Araport11 | Phytozome 13 | https://phytozome-next.jgi.doe.gov/ | <https://doi.org/10.1111/tpj.13415> |
| *Brachypodium distachyon* | v3.1 | Phytozome 13 | https://phytozome-next.jgi.doe.gov/ | <https://doi.org/10.1038/nature08747> |
| *Brassica rapa* | FPsc v1.3 | Phytozome 13 | https://phytozome-next.jgi.doe.gov/ |  |
| *Cardamine hirsuta* | CARH_v3.8 | *Cardamine hirsuta* Genetic and Genomic Resource | http://chi.mpipz.mpg.de/index.html | <https://doi.org/10.1038/nplants.2016.167> |
| *Citrus sinensis* | v1.1 | Phytozome 13 | https://phytozome-next.jgi.doe.gov/ | <https://doi.org/10.1038/nbt.2906> |
| *Cucumis sativus* | v1.0 | Phytozome 13 | https://phytozome-next.jgi.doe.gov/ |  |
| *Daucus carota* | v2.0 | Phytozome 13 | https://phytozome-next.jgi.doe.gov/ | <https://doi.org/10.1038/ng.3565> |
| *Diospyros lotus* | r1.1 | PersimmonDB | http://persimmon.kazusa.or.jp/index.html | <https://doi.org/10.1371/journal.pgen.1008566> |
| *Eucalyptus grandis* | v2.0 | Phytozome 13 | https://phytozome-next.jgi.doe.gov/ | <https://doi.org/10.1111/nph.13150> |
| *Gossypium raimondii* | v2.1 | Phytozome 13 | https://phytozome-next.jgi.doe.gov/ | <https://doi.org/10.1038/nature11798> |
| *Hordeum vulgare* | r1 | Phytozome 13 | https://phytozome-next.jgi.doe.gov/ | <https://doi.org/10.1038/sdata.2017.44> |
| *Lactuca sativa* | v8 | Phytozome 13 | https://phytozome-next.jgi.doe.gov/ | <https://doi.org/10.1038/ncomms14953> |
| *Medicago truncatula* | Mtv4.0v1 | Phytozome 13 | https://phytozome-next.jgi.doe.gov/ | <https://doi.org/10.1186/1471-2164-15-312> |
| *Mimulus guttatus* | v2.0 | Phytozome 13 | https://phytozome-next.jgi.doe.gov/ | <https://doi.org/10.1073/pnas.1319032110> |
| *Musa acuminata* | v1 | Phytozome 13 | https://phytozome-next.jgi.doe.gov/ | <https://doi.org/10.1038/nature11241> |
| *Nymphaea colorata* | v1.2 | Phytozome 13 | https://phytozome-next.jgi.doe.gov/ | <https://doi.org/10.1038/s41586-019-1852-5> |
| *Oropetium thomaeum* | v1.0 | Phytozome 13 | https://phytozome-next.jgi.doe.gov/ | <https://doi.org/10.1038/nature15714> |
| *Oryza sativa* | v7.0 | Phytozome 13 | https://phytozome-next.jgi.doe.gov/ | <https://doi.org/10.1093/nar/gkl976> |
| *Panicum hallii* | v3.2 | Phytozome 13 | https://phytozome-next.jgi.doe.gov/ | <https://doi.org/10.1038/s41467-018-07669-x> |
| *Panicum virgatum* | v5.1 | Phytozome 13 | https://phytozome-next.jgi.doe.gov/ | <https://doi.org/10.1038/s41586-020-03127-1> |
| *Physcomitium patens* | v3.3 | Phytozome 13 | https://phytozome-next.jgi.doe.gov/ | <https://doi.org/10.1111/tpj.13801> |
| *Populus trichopoda* | v4.1 | Phytozome 13 | https://phytozome-next.jgi.doe.gov/ | <https://doi.org/10.1126/science.1128691> |
| *Selaginella moellendorfii* | v1.0 | Phytozome 13 | https://phytozome-next.jgi.doe.gov/ | <https://doi.org/10.1126/science.1203810> |
| *Setaria viridis* | v2.1 | Phytozome 13 | https://phytozome-next.jgi.doe.gov/ | <https://doi.org/10.1038/s41587-020-0681-2> |
| *Solanum lycopersicum* | ITAG4.0 | Phytozome 13 | https://phytozome-next.jgi.doe.gov/ | <https://doi.org/10.1101/767764> |
| *Solanum tuberosum* | v6.1 | Phytozome 13 | https://phytozome-next.jgi.doe.gov/ | <https://doi.org/10.1093/gigascience/giaa100> |
| *Sorghum bicolor* | v3.1.1 | Phytozome 13 | https://phytozome-next.jgi.doe.gov/ | <https://doi.org/10.1111/tpj.13781> |
| *Triticum aestivum* | v2.2 | Phytozome 13 | https://phytozome-next.jgi.doe.gov/ | <https://doi.org/10.1126/science.1251788> |
| *Vitis vinifera* | v2.1 | Phytozome 13 | https://phytozome-next.jgi.doe.gov/ | <https://doi.org/10.1038/nature06148> |
| *Zea mays* | RefGen_v4 | Phytozome 13 | https://phytozome-next.jgi.doe.gov/ | <https://doi.org/10.1038/nature22971> |
