## Supplemental Table 3 for "Duplicate transcription factors *GT1* and *VRS1* regulate branching and fertile flower number in maize and *Brachypodium distachyon*"

**Supplementary Table 3: RNA-seq library statistics**

| **Sample ID** | **Genotype** | **Ear Size (cm)** | **Raw reads** | **Unique Mapped Reads** | **% Mapped** |
| --- | --- | --- | --- | --- | --- |
| jg107 | *vrl1* | 2.4 | 66513686 | 63192411 | 95.0% |
| jg121 | *wt* | 1 | 43535228 | 41369793 | 95.0% |
| jg145 | *gt1/+* | 1 | 49263064 | 46985881 | 95.4% |
| jg153 | *gt1;vrl1* | 0.5 | 49148840 | 46635295 | 94.9% |
| jg163 | *gt1/+* | 0.6 | 52906592 | 50502381 | 95.5% |
| jg164 | *wt* | 1.2 | 48005464 | 45617842 | 95.0% |
| jg166 | *gt1* | 0.8 | 67260448 | 64017557 | 95.2% |
| jg169 | *gt1/+;vrl1* | 1.2 | 54760834 | 52305387 | 95.5% |
| jg170 | *gt1;vrl1* | 1 | 63519594 | 60762018 | 95.7% |
| jg171 | *gt1* | 0.7 | 50993092 | 48688423 | 95.5% |
| jg175 | *gt1* | 1.5 | 54138178 | 51863869 | 95.8% |
| jg176 | *gt1/+;vrl1* | 0.6 | 47443100 | 45478770 | 95.9% |
| jg180 | *wt* | 1.5 | 48421500 | 46212929 | 95.4% |
| jg181 | *gt1/+;vrl1* | 1.6 | 63973078 | 60975702 | 95.3% |
| jg182 | *gt1* | 1.2 | 50499332 | 48210772 | 95.5% |
| jg183 | *gt1;vrl1* | 0.9 | 56852920 | 54354047 | 95.6% |
| jg187 | *wt* | 1 | 66351162 | 62996760 | 94.9% |
| jg188 | *gt1/+* | 0.6 | 74257912 | 70602437 | 95.1% |
| jg190 | *gt1;vrl1* | 0.7 | 55980966 | 52373908 | 93.6% |
| jg192 | *gt1* | 0.7 | 45785514 | 43766566 | 95.6% |
| jg193 | *gt1/+;vrl1* | 0.9 | 52484730 | 50218583 | 95.7% |
| jg195 | *gt1/+;vrl1/+* | 1.3 | 53972576 | 51583717 | 95.6% |
| jg196 | *gt1/+* | 1 | 47104762 | 44641938 | 94.8% |
| jg197 | *gt1* | 1.1 | 58182156 | 55681605 | 95.7% |
| jg198 | *vrl1* | 0.7 | 60831180 | 58653860 | 96.4% |
| jg199 | *gt1/+* | 1.5 | 54235196 | 51660781 | 95.3% |
| jg200 | *gt1/+;vrl1* | 1 | 63721640 | 60890462 | 95.6% |
| jg209 | *vrl1* | 1 | 57623028 | 55003631 | 95.5% |
| jg211 | *gt1/+;vrl1* | 1.8 | 46795426 | 45003612 | 96.2% |
| jg213 | *gt1/+;vrl1* | 0.7 | 48630918 | 46439020 | 95.5% |
| jg214 | *gt1/+;vrl1* | 0.9 | 49056470 | 46864733 | 95.5% |
| jg216 | *gt1/+;vrl1* | 2 | 45374430 | 43487084 | 95.8% |
| jg217 | *vrl1* | 0.7 | 64320370 | 62028279 | 96.4% |
| jg222 | *gt1/+* | 0.4 | 62829162 | 59440929 | 94.6% |
| jg223 | *gt1* | 0.3 | 62301002 | 59611975 | 95.7% |
| jg225 | *gt1* | 0.4 | 46650800 | 44349818 | 95.1% |
| jg228 | *gt1/+;vrl1* | 0.6 | 79175374 | 74884579 | 94.6% |
| jg231 | *gt1/+* | 0.8 | 74341142 | 70571970 | 94.9% |
| jg232 | *vrl1* | 0.4 | 58358920 | 53451357 | 91.6% |
| jg233 | *vrl1* | 0.7 | 60882652 | 57557331 | 94.5% |
| jg235 | *gt1/+;vrl1* | 0.7 | 76100510 | 67260392 | 88.4% |
| jg236 | *vrl1/+* | 0.9 | 48652514 | 46539832 | 95.7% |
| jg237 | *gt1/+;vrl1* | 0.8 | 61628424 | 58625660 | 95.1% |
| jg241 | *gt1/+;vrl1* | 0.4 | 50594696 | 47961878 | 94.8% |
| jg254 | *gt1;vrl1* | 0.8 | 53478724 | 50246787 | 94.0% |
| jg263 | *gt1;vrl1/+* | 0.8 | 57006136 | 54185489 | 95.1% |
| jg264 | *gt1;vrl1* | 0.5 | 67130492 | 63837083 | 95.1% |
| jg6 | *wt* | 1.7 | 45429992 | 43360307 | 95.4% |
| jg66 | *gt1;vrl1* | 2.3 | 50024284 | 48067385 | 96.1% |
| jg84 | *gt1/+;vrl1* | 1 | 47653964 | 45602640 | 95.7% |
| jg95 | *gt1;vrl1* | 1.6 | 60333108 | 57640096 | 95.5% |
| jg97 | *vrl1* | 1.3 | 51990478 | 49482610 | 95.2% |
| jg98 | *g1/+* | 1.8 | 48275476 | 46438862 | 96.2% |
